# So many genes, so little time: a practical approach to divergence-time estimation in the genomic era

**DOI:** 10.1101/114975

**Authors:** Stephen A. Smith, Joseph W. Brown, Joseph F. Walker

**Author notes:** Equal contribution (Joseph W. Brown, Joseph F. Walker). **Corresponding author:** Stephen A. Smith, Ecology and Evolutionary Biology, University of Michigan, Ann Arbor, Michigan, 48109, USA;.

## Abstract

Phylogenomic datasets have been successfully used to address questions involving evolutionary relationships, patterns of genome structure, signatures of selection, and gene and genome duplications. However, despite the recent explosion in genomic and transcriptomic data, the utility of these data sources for efficient divergence-time inference remains unexamined. Phylogenomic datasets pose two distinct problems for divergence-time estimation: (i) the volume of data makes inference of the entire dataset intractable, and (ii) the extent of underlying topological and rate heterogeneity across genes makes model mis-specification a real concern. “Gene shopping”, wherein a phylogenomic dataset is winnowed to a set of genes with desirable properties, represents an alternative approach that holds promise in alleviating these issues. We implemented an approach for phylogenomic datasets (available in **SortaDate**) that filters genes by three criteria: (i) clock-likeness, (ii) reasonable tree length (i.e., discernible information content), and (iii) least topological conflict with a focal species tree (presumed to have already been inferred). Such a winnowing procedure ensures that errors associated with model (both clock and topology) mis-specification are minimized, therefore reducing error in divergence-time estimation. We demonstrated the efficacy of this approach through simulation and applied it to published animal (Aves, Diplopoda, and Hymenoptera) and plant (carnivorous Caryophyllales, broad Caryophyllales, and Vitales) phylogenomic datasets. By quantifying rate heterogeneity across both genes and lineages we found that every empirical dataset examined included genes with clock-like, or nearly clock-like, behavior. Moreover, many datasets had genes that were clock-like, exhibited reasonable evolutionary rates, and were mostly compatible with the species tree. We identified overlap in age estimates when analyzing these filtered genes under strict clock and uncorrelated lognormal (UCLN) models. However, this overlap was often due to imprecise estimates from the UCLN model. We find that “gene shopping” can be an efficient approach to divergence-time inference for phylogenomic datasets that may otherwise be characterized by extensive gene tree heterogeneity.

## Introduction

Divergence-time estimation is a complicated, but often essential, step for many phylogenetic analyses. The sources of error include the ambiguous nature of fossil placement, model mis-specification (e.g., involving significant variation in the branchwise and/or sitewise rates of evolution), uncertainty in the phylogenetic tree, topological dissonance amongst gene trees due to incomplete lineage sorting, and complexity of the model for the molecular clock (e.g., Smith *et al*. 2010; Dornburg *et al*. 2012; Parham *et al*. 2012; Heath and Moore 2014; Beaulieu *et al*. 2015; Kumar and Hedges 2016). While fossils give the only available information for absolute age, their placement and age carry uncertainty. Multiple fossil calibrations and complicated tree shape priors can interact to further complicate molecular dating (Zhu *et al*. 2015; Heled and Drummond 2015; Rannala 2016; dos Reis 2016; Brown and Smith 2017). Rate variation is common among individual branches of a phylogeny and can constitute extensive deviations from the molecular clock. As a result, complex models have been developed to accommodate for these deviations (Sanderson 2002; Drummond *et al*. 2006; Drummond and Suchard 2010). However, these more parameter-rich models also carry with them significant uncertainty and can, when the data deviate significantly from the model, lead to biased results (e.g., Worobey *et al*. 2014). Despite these difficulties, researchers continue to use divergence-time estimates extensively as they remain essential for many downstream evolutionary and comparative analyses.

Datasets based on thousands of genes from genomes and transcriptomes have emerged as a major tool in addressing broad evolutionary questions including, but not limited to, phylogenetic reconstruction, gene and genome duplication, and inference of molecular evolutionary patterns and processes. And while these datasets have been used for divergence-time estimation (e.g., Jarvis *et al*. 2014b; Prum *et al*. 2015), their overall utility for divergence-time analyses has not been fully examined. In particular, it is unclear whether within these enormous datasets there exist nearly clock-like gene regions that may aid in producing lower error divergence-time estimates. While some authors of recent large genomic analyses, such as Jarvis *et al*. (2014b), have suggested restricting analysis to clock-like genes, a repeatable and fast procedure to identify these genes has not been explored for phylogenomics, and an examination of the frequency of these genes in empirical datasets has not yet been conducted.

Researchers can take steps to ease sources of errors for divergence-time analyses. For example, better use of fossils in temporal calibrations can dramatically improve estimations (e.g., Parham *et al*. 2012; Ksepka *et al*. 2015), as does better accounting for rate variation in the molecular models by improving model fit. Several relaxed clock models have been introduced over the last few decades to accommodate rate heterogeneity because most data do not conform to a strict clock. The most commonly used relaxed clock methods include penalized likelihood (PL, Sanderson 2002) as implemented in **r8s** (Sanderson 2003) (and more recently in **treePL**; Smith and O’Meara (2012)), and Bayesian uncorrelated rate models (e.g., the uncorrelated lognormal (UCLN) model; Drummond *et al*. (2006)) as implemented in **BEAST** (Drummond and Rambaut 2007) and **MrBayes** (Ronquist *et al*. 2012), although many other methods have been developed and new ones are continually released (e.g., Takezaki *et al*. 1995; Thorne and Kishino 2002; Lartillot and Philippe 2004; Britton *et al*. 2007; Lepage *et al*. 2007; Rannala and Yang 2007; Drummond and Suchard 2010; Tamura *et al*. 2012; Heath *et al*. 2014; Ronquist *et al*. 2016; Lartillot *et al*. 2016). The diversity of techniques is matched with a variety of different inputs. For example, PL implementations minimally require an estimated phylogram, calibration, smoothing penalty value, and alignment size, while full Bayesian methods minimally require an alignment and priors to be set for each parameter, including any fossil calibrations.

Bayesian methods that use relaxed clock models, such as those implemented in **BEAST**, **MrBayes**, and **PhyloBayes** (Lartillot and Philippe 2004), simultaneously estimate phylogenetic relationships and divergence-times, and so may be preferred over other approaches as Bayesian methods incorporate uncertainty more easily and explicitly. However, the computational burden of these simultaneous reconstruction methods limit their use to smaller datasets (i.e., excluding entire genomes and transcriptomes). Fortunately, a prescient solution to this dilemma was proffered two decades ago with the concept of “gene shopping” (Hedges *et al*. 1996; Kumar and Hedges 1998), wherein available genes are filtered by how well they conform to a molecular clock. [A related procedure, “taxon-shopping” (Takezaki *et al*. 1995; van Tuinen and Hedges 2001; van Tuinen and Dyke 2004), prunes taxa from an alignment until the dataset no longer rejects a molecular clock test. We do not consider this approach here.]. Using “gene shopping”, it should be possible to reduce larger datasets to alignments that are capable of being analyzed by Bayesian methods. However, despite the having been available for decades, “gene shopping” has not been widely applicable before the recent development of next-generation sequencing techniques because of the relatively small number of genes available for any single clade (outside of model organisms). However, as genomic and transcriptomic datasets have become more readily available, “gene shopping” holds tremendous promise as a tool for inferring phylogenetic timescales. Nevertheless, the utility of these large genomic datasets for divergence-time estimation and the distribution of lineage-specific rate heterogeneity has yet to be fully explored.

Next generation sequencing techniques have dramatically increased the number of gene regions available for phylogenetic analysis. This has stimulated research into questions that are specifically pertinent to datasets with hundreds or thousands of genes. What is the best method for reconstructing the species tree (e.g., Gatesy and Springer 2014; Mirarab *et al*. 2014; Roch and Warnow 2015)? How many genes support the dominant species tree signal (e.g., Salichos *et al*. 2014; Smith *et al*. 2015)? Genomic datasets also allow us to examine the extent of molecular rate variation in genes, genomes, and lineages. For example, Yang *et al*. (2015) explored the distribution of lineage-specific rate heterogeneity throughout transcriptomes of the plant clade Caryophyllales as it relates to life history. Jarvis *et al*. (2014b), analyzing a genomic dataset of birds, explored rate heterogeneity and selection as it relates to errors in phylogeny reconstruction in a genomic dataset of birds. Recently, the clock-likeness of phylogenomic datasets has come of interest to the community. For example, Doyle *et al*. (2015) attempt to identify strictly clock-like genes in order to avoid long branch attraction artifacts in phylogenetic inference. Jarvis *et al*. (2014b) filtered gene regions by inferred mean coefficient of rate variation (a measure of clock-likeness) from full Bayesian analyses of each gene, to identify “clock-like” genes explicitly for divergence-time estimation. However, while these authors have conducted filtering analysis on their genomic data, a thorough examination of lineage-specific rate heterogeneity across clades for divergence-time estimation has not been conducted.

Here we present one means of utilizing genomic data for estimating divergence-times by introducing a simple winnowing procedure to identify informative and low variance genes (see description below). Genes that satisfy these criteria may enable convenient and efficient divergence-time estimation for large datasets as methods such as clock partitioning are difficult when dealing with dozens, hundreds, or even thousands of genes Duchêne *et al*. (2013). Additionally, this procedure can be used to examine the overall distribution of evolutionary rate heterogeneity, bipartition concordance, and potential utility of genes for divergence-time analysis. It is assumed that a researcher will possess a species tree in order to determine what gene regions best conform to that hypothesis. This can be especially important given the underlying conflict present in most genomic datasets. The procedure holds promise that, while various relaxed clock models are available (conducive to accommodating different forms of rate heterogeneity), data have lower variance and may not require as extensive rate modeling will enable fast, accurate, and precise divergence-time estimates (see also To *et al*. 2015). Finally, we examine six genomic and transcriptomic datasets across animals and plants and with different temporal and taxonomic scopes to examine the extent of lineage-specific rate heterogeneity. We investigate the distribution of variation in the branchwise rates of evolution across thousands of genes to understand whether these new genomic resources may improve divergence-time estimation by enabling analysis using simpler models of molecular evolution.

## Materials and Methods

### Description of procedures

Because phylogenomic datasets consist of hundreds to thousands of genes, we developed a winnowing procedure intended to mimic a “gene shopping” exercise (see Fig. 1). Our approach has two requirements: (i) a species tree topology must be provided (presumed to have been inferred by some means), and (ii) individual gene trees must be rooted (this is typically performed using outgroups). The filtering procedure itself relies on three criteria relevant to divergence-time estimation: (i) a root-to-tip variance statistic (an indication of clock-likeness), (ii) a bipartition calculation to determine the similarity to the provided species tree, and (iii) the total tree length (i.e., discernible information content). Our implementation of this pipeline sorts first by the similarity to the species tree, then root-to-tip variance, and finally tree length. This procedure produces a ranked list of genes that jointly satisfy the filtering criteria. Below we demonstrate the efficacy of this approach using both simulated and empirical data. The analyses performed herein can be conducted using the **SortaDate** package (with source code and instructions available at https://github.com/FePhyFoFum/sortadate). This package is written in Python and available as an open source set of procedures. In some cases, external programs are used (e.g., those found in the **Phyx** package; Brown *et al*. (2017)) that are also open source and freely available.

**Figure 1:**
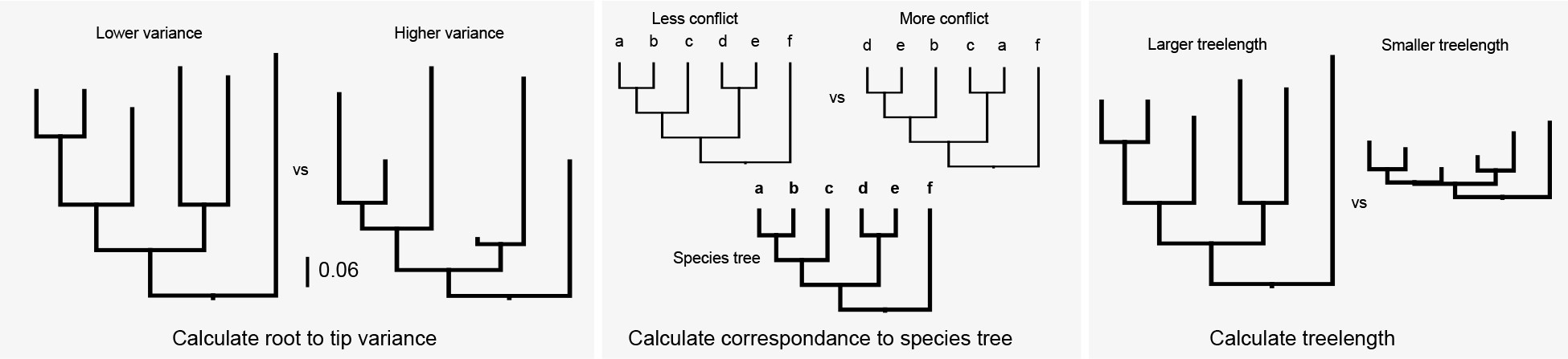
Winnowing criteria used for sorting genes for use in divergence-time inference analyses. The order presented here is arbitrary.

### Dataset processing

We analyzed six published phylogenomic datasets to explore rate and topological heterogeneity: birds (Aves) (BIR, Jarvis *et al*. 2014b), carnivorous Caryophyllales (CAR, Walker *et al*. 2017), the broader Caryophyllales (CARY, Yang *et al*. 2015), Vitales (VIT, Wen *et al*. 2013), Hymenoptera (HYM, Johnson *et al*. 2013), and millipedes (Diplopoda) (MIL, Brewer and Bond 2013). The range in datasets spans different taxonomic groups (e.g., animals vs. plants), datasets sizes (e.g., CAR vs CARY), and age (e.g., from hundreds of millions of years to within the last hundred million years). For all datasets except BIR orthologs were identified using the Maximum Inclusion method of Yang and Smith (2014). These data are available at https://github.com/FePhyFoFum/SDMIData. For each ortholog we have an alignment and maximum likelihood (ML) gene tree inferred in **RAxML** v8.2.3 (Stamatakis 2014) using the GTR+G model for DNA and WAG+G for amino acids. For the BIR dataset we used the exon alignments and inferred phylograms available online (Jarvis *et al*. 2014a; http://gigadb.org/dataset/101041). Gene trees, regardless of the source of orthologs, were then rooted and SH-like tests were performed to assess confidence in edges (Anisimova and Gascuel 2006).

### Gene tree analyses

Because deviation from the clock is empirically manifest in a phylogram as a departure from ultrametricity, we measured the variance of root-to-tip lengths for each ML gene tree phylogram. This was performed on each rooted ortholog, for which outgroups were removed, with the **pxlstr** program of **Phyx** package (Brown *et al*. 2017). We performed the standard clock test for each ortholog (Muse and Weir 1992), with outgroup removed, using **PAUP\*** v4.0a151 (Swofford 2001) by calculating the ML score for a gene both with and without assuming a clock, and then performing a likelihood ratio test. In addition to assessing the clock-likeness of genes, we also compared gene tree topologies to the corresponding published species tree topology. Branch lengths were not available for some species trees. To compare the individual gene trees to their corresponding species trees, we conducted bipartition comparison analyses on each gene tree using **pxbp** from the **Phyx** package (procedure described in Smith *et al*. 2015).

### Simulations

We conducted simulations to examine expectations of rate variation given clock-like, noisy clock-like, and uncorrelated lognormal data. We first generated simulated clock-like data using **Indelible** v1.03 (Fletcher and Yang 2009) using the WAG model with 500 characters for amino acid datasets, and JC with 1500 characters for nucleotide datasets, on each of the empirical species tree topologies. For these data simulations, because the species tree often had no branch lengths available, node heights were first simulated randomly using **Indelible** and then the tree was rescaled to a height of 0.25, 0.5, or 0.75. We used the trees generated by **Indelible** to further simulate 100 noisy clock (rate=1.0, noise=0.25, and rate=1.0, noise=0.75) and uncorrelated lognormal (UCLN) trees (mean.log=-0.5, stdev.log=0.5, and mean.log=-0.5, stdev.log=1.0) using **NELSI** v0.21 (Ho *et al*. 2015). We note the ‘noise’ in **NELSI** corresponds to the standard deviation of a normal distribution with mean = 0. For the noisy clock, branch-specific rates are a sum of the global rate (here, 1.0) and a draw from this normal distribution. The simulations with noise=0.75 thus are only loosely clock-like, and serve as a comparison between the more clock-like (noise=0.25) and UCLN analyses. We used **RAxML** to reconstruct each of these datasets. For each simulation, we examined the rate variation and the root-to-tip length variation on the reconstructed phylograms.

While the focus of this study is not the performance of divergence-time estimation methods, we nevertheless examine an exemplar from the simulations to ascertain the variation in the results given different clock models. We used one random realization of node heights as simulated from the **Indelible** analyses as described above to generate two datasets with **NELSI**. One dataset had three genes generated from a clock rate of 1 and noise at 0.25, and the other dataset had three genes generated from a UCLN model and mean.log at −0.5 and sd.log=1. As above, each amino acid gene consisted of 500 residues, while DNA genes consisted of 1500 nucleotides. For each simulation, all three genes shared a common topology (but with different edge lengths, as our filtering procedure involves a single focal topology). For both the noisy clock-like and UCLN datasets, we conducted **BEAST** analyses with both a clock model and a UCLN model. Thus we have two scenarios where the generating and inference clock models are identical, and two where they are mismatched. A birth-death tree prior was used as the prior for all node heights, and runs were conducted for 10 million generations with the first 10% discarded as burnin. Results were summarized using **treeannotator** from the **BEAST** package. Median node heights as well as 95% highest posterior density (HPD) node heights were compared between the simulated datasets and the tree used to generate these datasets.

Finally, to demonstrate the inference precision afforded by our winnowing procedure, we conducted simulations where heterogeneity was restricted to overall molecular substitution rates (i.e., tree lengths) and the degree of unclock-like noise involved. In all, 1000 Yule trees with a speciation rate of 1.0 were simulated for 12 taxa using **pxbdsim** from the **Phyx** package. For each tree, we simulated 1000 nucleotide alignments under JC, to reduce additional variation, and a strict clock model with varying degrees of noise. ML trees for each alignment were inferred using **RAxML**. From each pool of alignments we constructed 2 datasets: (i) three genes chosen randomly, and (ii) three genes selected from the **SortaDate** procedure. All datasets were analyzed in **BEAST** under both strict clock and UCLN models, where genes shared a clock model but had gene-specific relative rates and individual GTR+G substitution models. The tree topology was fixed to match the true tree. As above, analyses were conducted for 10 million generations with the first 10% discarded as burnin. Finally, we compare the inferred median node ages from the resulting maximum clade credibility trees to the true simulated tree. We note that because within a simulation there is no real topological conflict, any heterogeneity involved in inference accuracy involves (i) the gene selection criteria and (ii) the clock model used for inference.

### Sorting and dating analyses on real data

In addition to these analyses on simulated datasets, we conducted divergence-time analyses on a subset of the empirical datasets. For these examples we limited the results to the top three genes reported from the sorting procedure, although for empirical analyses one should choose the number of genes by carefully examining the filtering results. Because we filtered for genes that were consistent with the species tree, these genes were then concatenated and the topology was fixed to be consistent with the species tree. We applied individual substitution models to each gene within a dataset. However, given that the genes were filtered both to match the species tree and for clock-likeness, we modeled all genes with a single molecular clock (albeit with gene-specific relative rates). For each dataset, we conducted two **BEAST** analyses, one assuming a strict clock and the other assuming a UCLN model. Because specific dates were not the focus of this examination, the birth-death tree prior was used instead of fossil priors for nodes. The analyses were run in duplicate for 50 million generations with the first 10% discarded as burnin. Replicate MCMC logs were concatenated while removing burnin using the **pxlog** program from the **Phyx** package, and finally summarized using **treeannotator** as above. Median node heights and 95% HPD node heights were compared between the clock and UCLN runs as the node heights on the true phylogeny are unknown.

## Results and Discussion

A fundamental question for each empirical dataset examined here is whether there are clock-like gene regions present within the genome? Results were varied, from 0.4% of genes passing the strict clock test for the VIT dataset to 17% for the MIL dataset (see Table 1). Those datasets with fewer clock-like genes may have underlying models that deviate significantly from the clock. However, there are several reasons that, even with a strict clock model underlying these data, we might expect deviations from a clock. For example, we expect stochastic differences to accumulate with increased sampling (i.e., more edges) and older trees (i.e., longer edges). Within the datasets examined here, two, the CAR and CARY, overlap in sampling with the CAR dataset having far fewer taxa in total. The broader sampling within the CARY dataset, in addition to greater dataset size, also results in including more known life-history shifts (Yang *et al*. 2015). These differences may account for the variation between these two datasets with the CARY dataset having significantly more variation and fewer clock-like genes. As for clade age, HYM and MIL datasets include divergence-times that are significantly older than the other datasets, which may account for the rate variation within those data. Nevertheless, each dataset had at least a few orthologs that passed a strict clock test even if these orthologs were in the small minority.

**Table 1:** Dataset properties and results of likelihood ratio tests for strict clock-like gene behaviour.

| Dataset | Ingroup | Data type | Orthologs | Clocklike (%) |  |
| --- | --- | --- | --- | --- | --- |
| BIR | 46 | DNA | 7116 | 440 | (6.18) |
| CAR | 11 | AA | 3767 | 274 | (7.27) |
| CARY | 68 | AA | 583 | 3 | (0.51) |
| HYM | 18 | AA | 1161 | 22 | (1.89) |
| MIL | 10 | AA | 152 | 26 | (17.10) |
| VIT | 16 | DNA | 2267 | 8 | (0.35) |

Because passing a clock test does not necessarily indicate that the gene would be good for phylogenetic reconstruction (e.g., if the rate of evolution is so slow as to be uninformative), we also measured tree length and root-to-tip variance for each ortholog (see Figs. 2, 3). Clock tests are stringent in their need to conform to the clock (see below) and so by examining the root-to-tip variation and lineage-specific variation, we are more directly examining the deviation from ultrametricity. Although this is primarily descriptive and does not include a formal test, this provides an easily interpretable characterization of rate variation. We found that the datasets varied dramatically with no discernible general pattern for both root-to-tip variance and tree length. For example, the BIR dataset demonstrated very little molecular evolution as demonstrated by the short tree lengths. For this dataset, we analyzed nucleotides (rather than amino acids) to maximize tree lengths as Jarvis *et al*. (2014b) demonstrated low rates of evolution, especially deep in the phylogeny. However, the inferred rates of evolution (as determined by overall tree length) were still low. Given the difficulty in resolving the avian phylogeny, this pattern is, perhaps, to be expected (Jarvis *et al*. 2014b). This same pattern is present in the VIT dataset, though this was not explored as thoroughly in the original publication. Both the CAR and CARY datasets showed a pattern of increasing variance with greater tree length (Fig. 2). This contrasts with the HYM and MIL datasets that were clock-like even with longer tree lengths (Fig. 3). Lineage-specific rate variation in each dataset was idiosyncratic with most extreme variation in the outgroups. While outgroups were excluded for clock tests and in determining root-to-tip variance for “gene shopping”, we allowed outgroups to remain for lineage-specific rate variation analyses as in the right hand plots of Fig. 3. The VIT dataset was an exception with several lineages other than the outgroup having high rates. In each dataset, there were genes that fell within the distribution of simulated trees that are clock-like or clock-like with low noise.

**Figure 2:**
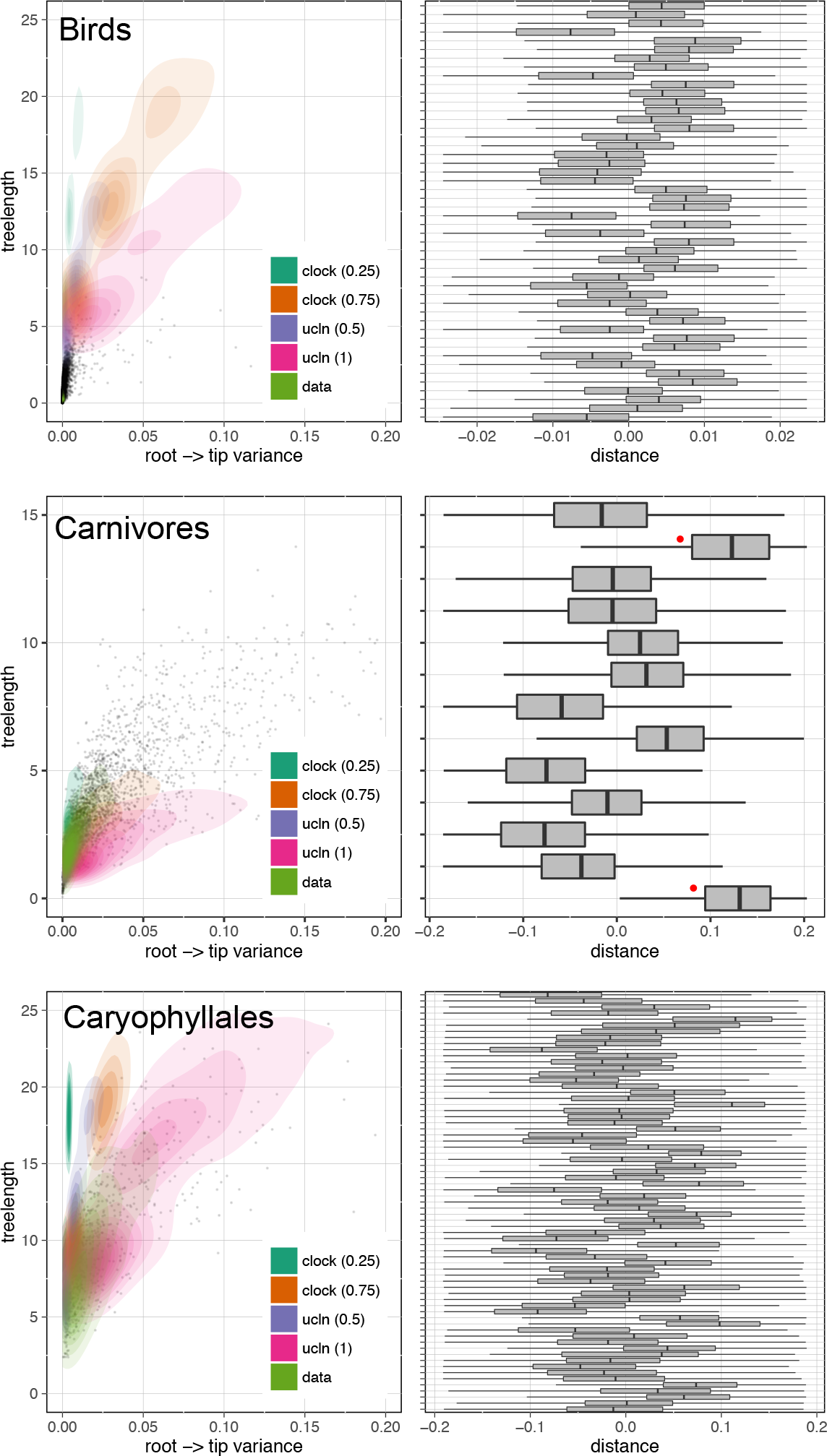
Gene tree properties for the BIR, CAR, and CARY datasets. Left: relationship between root-to-tip variance and tree length for simulated (*clock* and *ucln*) and empirical (*data*) datasets). Each simulation condition consists of 100 simulated datasets. Contours represent densities, while grey dots represent raw empirical values. Right: tip-specific root-to-tip variance for empirical datasets. Here, 0 represents the mean root-to-tip across all genes and all lineages. Red dots indicate outgroup taxa.

**Figure 3:**
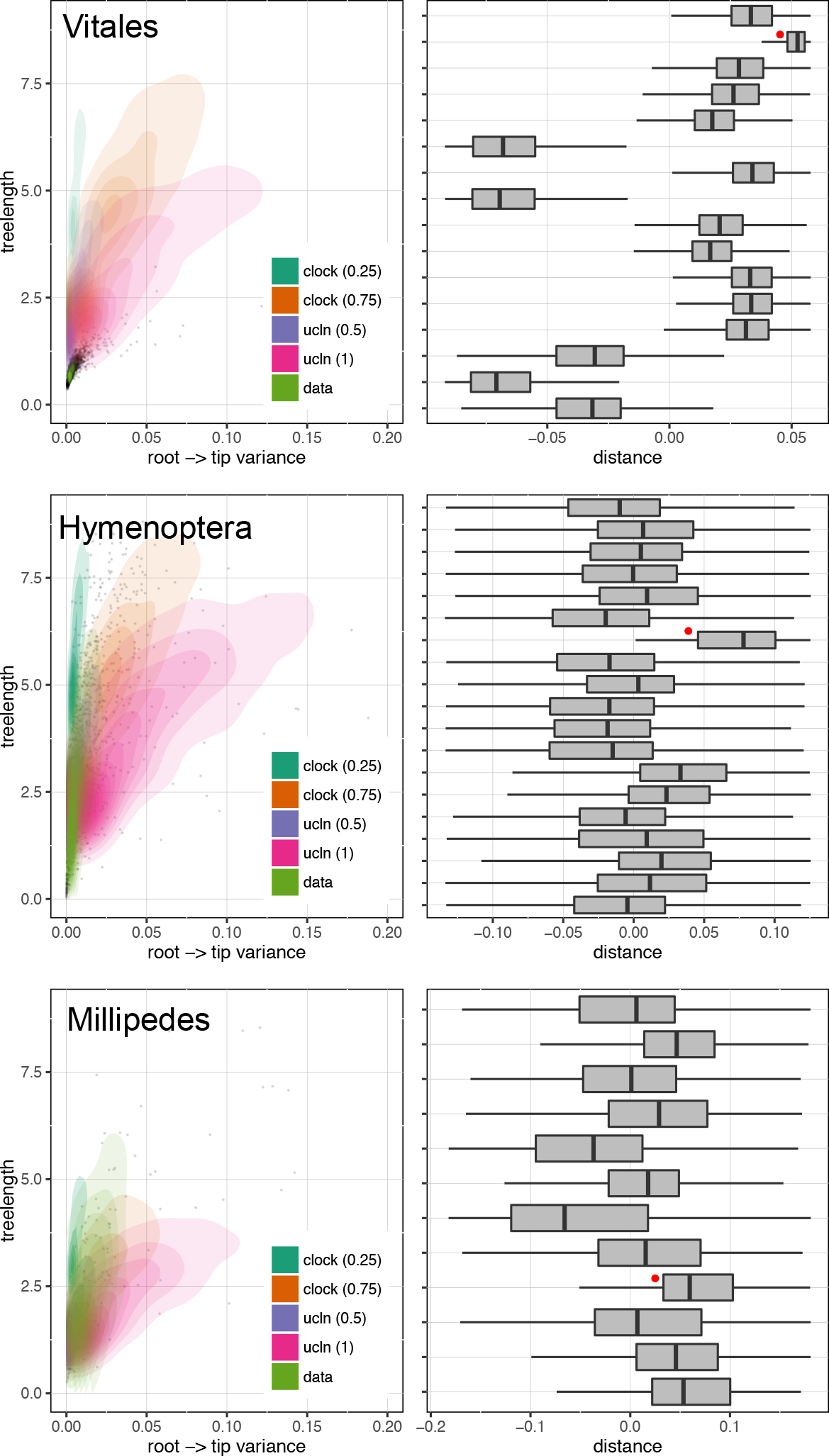
Gene tree properties for the VIT, HYM, and MIL datasets. See Fig. 2 for a description of the plotting attributes.

One potential benefit of identifying orthologs with lower lineage-specific rate variation within phylogenomic datasets is to use these, or a subset of these, orthologs to conduct divergence-time analyses. The hope is that by using clock-like genes, we may overcome or alleviate the impact of lineage-specific rate variation on the error of divergence-time analyses. The non-identifiability of rates and dates (e.g., longer branch lengths may be the result of a long time or fast evolution) is exacerbated by lineage-specific rate heterogeneity. We used a subset of orthologs to conduct divergence-time analyses and we implemented a sorting procedure (available in **SortaDate**) to (i) filter the genes that best reflect the species tree (i.e., higher bipartition concordance with the species tree), (ii) have lower root-to-tip variance (i.e., most clock-like), and (iii) discernible amounts of molecular evolution (i.e., greater tree length; Fig. 1). For each empirical dataset, we generated such an alignment (see Table 2). The genes that were filtered and used for divergence-time analyses for the BIR, CARY, VIT, and HYM datasets rejected the clock. The genes for the CAR and MIL datasets either didn’t or weakly rejected the clock. Resulting HPD trees were rescaled so that the root heights were equivalent to allow for easier comparisons between datasets. Typically, fossil placements would be used for scaling but because these are not intended to be runs for future use, we eliminated fossil placements as one source of variation. We found rough correspondence of node heights between the clock and UCLN analyses, especially for the four smallest datasets (see Fig. 4). The UCLN analyses, as expected, had far greater variance in the 95% HPDs for node ages. We found the greatest differences in the larger BIR and CARY datasets (see Table 3) where there are major differences in tree heights. This may reflect the size of the dataset or the underlying rate variation in the datasets. In general, strict clock estimates resulted in younger median node ages than analogous UCLN estimates, as well as younger maximum and older minimum 95% HPD values (see Table 3). The coefficient of rate variation statistics (an measure of clock-likeness) for UCLN runs ranged from the lowest mean values of 0.2358 (stErr=0.0347) in HYM to the highest of 1.2464 (stErr=0.1135) in CARY.

**Table 2:** Properties of the filtered genes used in the empirical dating analyses. Variance regards the root-to-tip paths. Tree length is measured in units of expected substitutions per site across all branches. Bipartition proportion measures agreement to the species tree topology (1.0 indicates complete concordance).

| Gene name | Variance | Tree length | Bipartition proportion |
| --- | --- | --- | --- |
| **BIR** |  |  |  |
| 12969 | 0.000791644 | 2.73068 | 0.5 |
| 1173 | 0.00205589 | 3.01712 | 0.457 |
| 12123 | 8.32228e-05 | 0.825943 | 0.413 |
| **CAR** |  |  |  |
| cluster259MIortho7 | 9.07832e-05 | 0.346618 | 1.0 |
| cluster3790MIortho1 | 0.000245644 | 0.739886 | 1.0 |
| cluster234MIortho1 | 0.0004849 | 1.19575 | 1.0 |
| **CARY** |  |  |  |
| cc7674-1-1to1ortho | 0.0183029 | 10.9821 | 0.701 |
| cc4427-1MIortho1 | 0.0093838 | 8.7827 | 0.657 |
| cc7873-1MIortho1 | 0.0206222 | 10.4773 | 0.657 |
| **HYM** |  |  |  |
| cluster3024-1-1ortho1 | 0.00159156 | 2.64137 | 0.706 |
| cluster5160-1-1ortho1 | 0.00294197 | 2.0815 | 0.706 |
| cluster1251-1-1ortho1 | 0.00621115 | 4.99913 | 0.706 |
| **MIL** |  |  |  |
| cluster89-1-1ortho1 | 0.00200945 | 0.909593 | 0.875 |
| cluster1437-1-1ortho1 | 0.00872612 | 2.96511 | 0.875 |
| cluster1615-1-1ortho1 | 0.010942 | 3.56434 | 0.875 |
| **VIT** |  |  |  |
| cluster9579-1MIortho1 | 0.000978163 | 0.519373 | 1.0 |
| cluster1236-1MIortho1 | 0.00106778 | 0.547562 | 1.0 |
| cluster461-1MIortho1 | 0.001227 | 0.607536 | 1.0 |

**Table 3:** The cumulative difference across all nodes in the height, lower 95% HPD, and higher 95% HPD of each node comparing the UCLN estimates to the strict clock estimates from the individual empirical dating analyses. A value lower than 0 results when the cumulative difference in the clock values of height or HPD are younger than the associated UCLN values.

| Dataset | Height | Lower | Higher |
| --- | --- | --- | --- |
| **BIR** | -0.26 | 0.27 | -1.49 |
| **CAR** | -0.004 | 0.04 | -0.2 |
| **CARY** | -3.93 | 0.52 | -8.56 |
| **HYM** | -0.12 | 0.1 | -0.63 |
| **MIL** | -0.09 | 0.04 | -1.12 |
| **VIT** | -0.02 | 0.08 | -0.56 |

**Figure 4:**
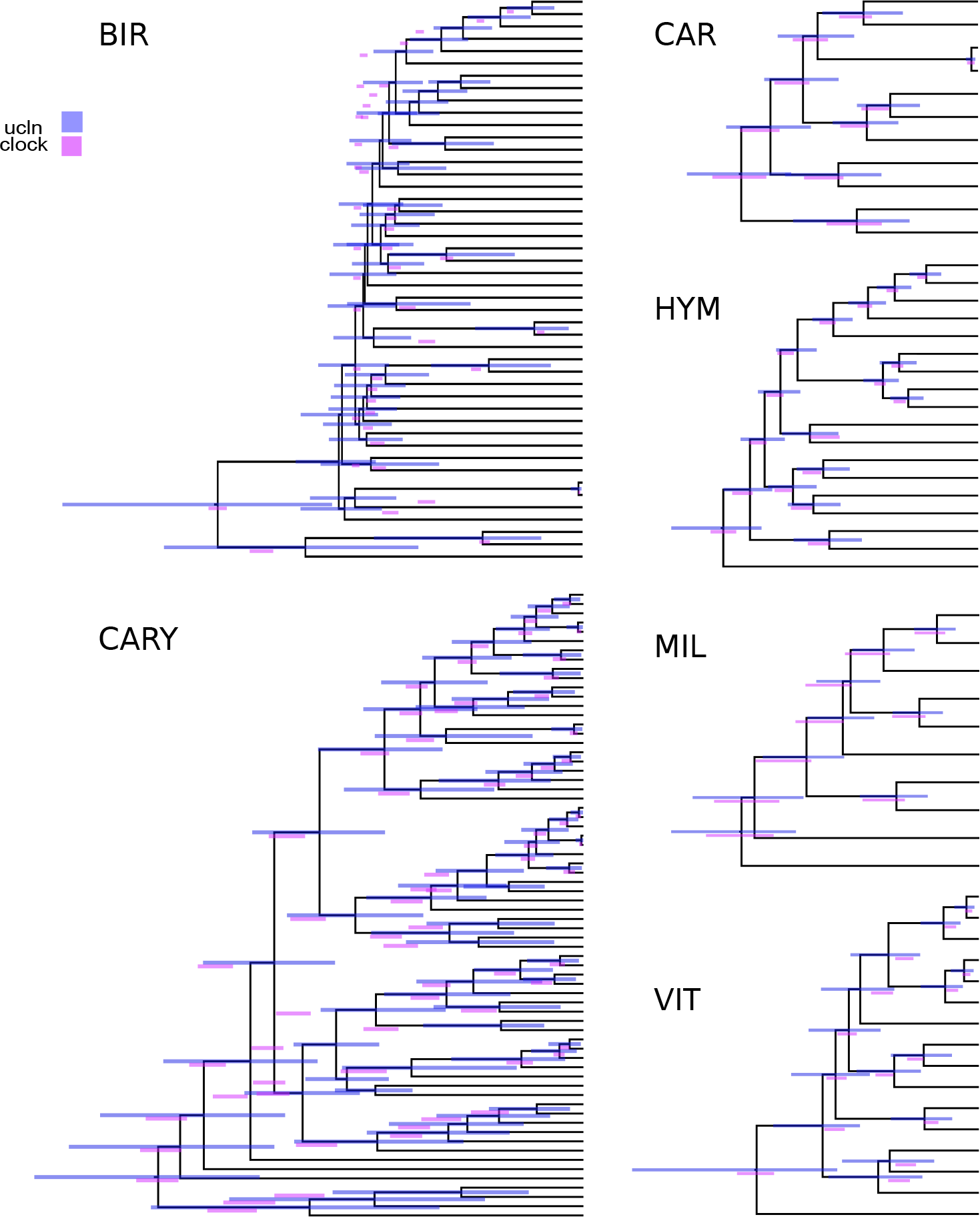
A comparison of strict clock and UCLN estimates of node ages for the six curated empirical datasets. Bars represent 95% HPD intervals and overlay the UCLN maximum clade credibility tree.

As is always the problem with real datasets, the true divergence-times are unknown. So we conducted exemplar analyses. For each empirical dataset, we simulated data for three genes under both noisy clock and UCLN models to examine the variation in the resulting divergence-time analyses where the true dates were known. For these simulated datasets, a strict clock was rejected in each case, including those datasets that were simulated under a clock with noise. We compared the resulting node heights from the divergence-time analyses under clock and UCLN models with the tree used for simulation (see Tables 4, 5 and Fig. 5). For the datasets generated under a noisy clock model, more of the true node heights were found in the 95% HPD interval when using the UCLN model for inference than the strict clock model for inference. However, the precision as measured by the total width of the 95% HPD interval for the UCLN runs were much lower than the clock runs (see Tables 4, 5). Those nodes that were not within the interval of the 95% HPD when using the strict clock model for reconstruction, were close to the true value. So, while fewer true node ages were contained in the strict clock HPDs, the overall error rate was lower. For example, in the CARY dataset, while fewer nodes in the clock estimate were found to be within the interval (52 vs 67 for the UCLN), the distance of the interval from the estimate was lower for the clock dataset for both the high and low value for the 95% HPD. Stated another way, the UCLN intervals were large enough that the true age was often included, but this was at the cost of far lower precision. Because of this error relative to the strict clock, the UCLN perhaps should not be the preferred model, especially if the researcher is going to use a single summary tree for future analyses.

**Table 4:** Assessment of dating error for the clock (CL) and UCLN (UC) analyses of the simulated *clock* data. All measures involve distance from the true node age, and are cumulative sums across all nodes. Height is the inferred node age. Lower and Higher regard the 95% HPD node age bounds. Nodes indicates the number of true node ages contained within the HPD interval. Error is the total error involved, equivalent to Low + High.

| Dataset | Height |  | Lower |  | Higher |  | Nodes |  | Error |  |
| --- | --- | --- | --- | --- | --- | --- | --- | --- | --- | --- |
|  | CL | UC | CL | UC | CL | UC | CL | UC | CL | UC |
| **BIR** | 0.63 | 0.5 | 0.48 | 0.69 | 0.96 | 1.33 | 16 | 39 | 1.44 | 2.02 |
| **CAR** | 0.26 | 0.2 | 0.37 | 0.85 | 0.25 | 0.63 | 5 | 12 | 0.62 | 1.49 |
| **CARY** | 0.54 | 0.63 | 1.23 | 2.12 | 0.64 | 1.11 | 52 | 67 | 1.88 | 3.24 |
| **HYM** | 0.16 | 0.76 | 0.21 | 0.65 | 0.38 | 0.94 | 15 | 3 | 0.58 | 1.59 |
| **MIL** | 0.17 | 0.42 | 0.12 | 0.45 | 0.33 | 0.46 | 5 | 3 | 0.44 | 0.91 |
| **VIT** | 0.27 | 0.26 | 0.42 | 0.32 | 0.2 | 0.27 | 5 | 8 | 0.62 | 0.59 |

**Table 5:** Assessment of dating error for the clock (CL) and UCLN (UC) analyses of the simulated *ucln* data. All measures involve distance from the true node age, and are cumulative sums across all nodes. Height is the inferred node age. Lower and Higher regard the 95% HPD node age bounds. Nodes indicates the number of true node ages contained within the HPD interval. Error is the total error involved, equivalent to Low + High.

| Dataset | Height |  | Lower |  | Higher |  | Nodes |  | Error |  |
| --- | --- | --- | --- | --- | --- | --- | --- | --- | --- | --- |
|  | CL | UC | CL | UC | CL | UC | CL | UC | CL | UC |
| **BIR** | 1.26 | 3.21 | 1.26 | 6.43 | 1.37 | 1.52 | 12 | 24 | 2.64 | 7.95 |
| **CAR** | 0.76 | 0.69 | 0.89 | 1.59 | 0.68 | 0.58 | 2 | 9 | 1.57 | 2.17 |
| **CARY** | 2.29 | 3.51 | 2.37 | 8.98 | 2.38 | 4.97 | 15 | 55 | 4.75 | 13.95 |
| **HYM** | 0.14 | 0.91 | 0.61 | 3.01 | 0.61 | 1.58 | 18 | 16 | 1.22 | 4.65 |
| **MIL** | 0.14 | 0.61 | 0.32 | 1.6 | 0.57 | 1.17 | 11 | 10 | 0.89 | 2.77 |
| **VIT** | 0.29 | 1.12 | 0.82 | 2.43 | 0.29 | 0.73 | 14 | 9 | 1.11 | 3.16 |

**Figure 5:**
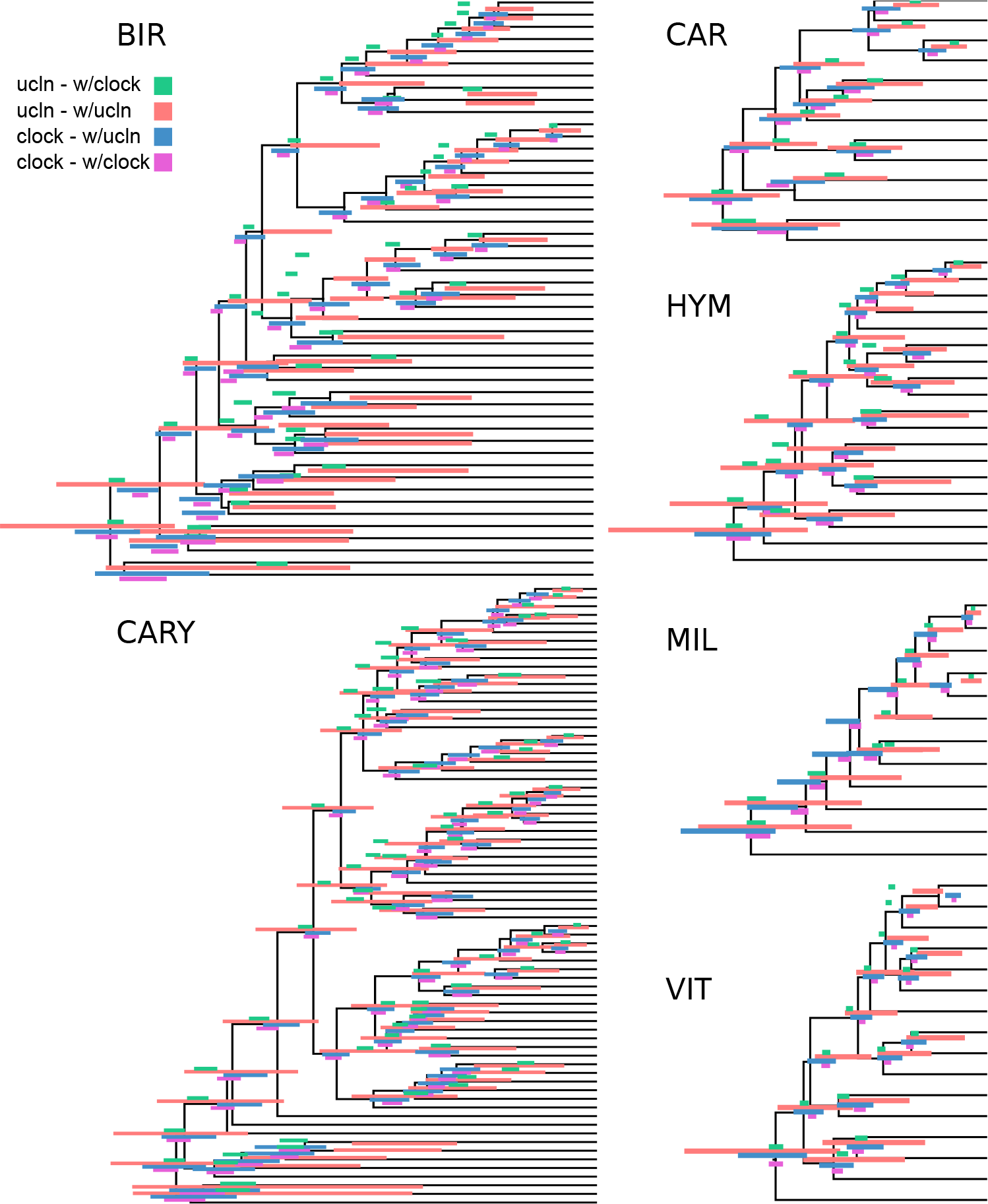
A comparison of strict clock and UCLN estimates of node ages for the simulated *clock* and *ucln* datasets. Each simulation condition consists of three genes. Red and pink are scenarios where the generating and inference models are identical, while green and blue are where the models are mismatched. Bars represent 95% HPD intervals and overlay the true simulated tree.

Our results on the empirical datasets, both with simulated and real data, suggest that a ‘gene shopping’ approach is can yield more precise estimates of divergence times. However, because the complexity of the underlying gene trees for these empirical datasets can complicate our interpretation of divergence-time results, we conducted additional simulations where the underlying true trees and true divergence-times were known. In each simulation, genes shared the same underlying species tree but varied in the rate and noise of the underlying clock model. In comparing the divergence-times estimated using the genes identified from SortaDate versus those chosen at random, it is evident that winnowed genes produced more precise estimates ((Fig. 6). This is expected as SortaDate aims to reduce variation in aspects of genetic data that increase variance. In addition, reducing inference model complexity to a strict clock reduced error substantially. Given that the data were generated with a clock, this was not unexpected but we note that the UCLN model did not perform like a clock when the data are essentially clock-like. Therefore, when possible, it seems prudent to restrict model complexity (i.e., when the data conform to the model). However, whether this is possible will depend entirely on the properties of the empirical dataset in question.

**Figure 6:**
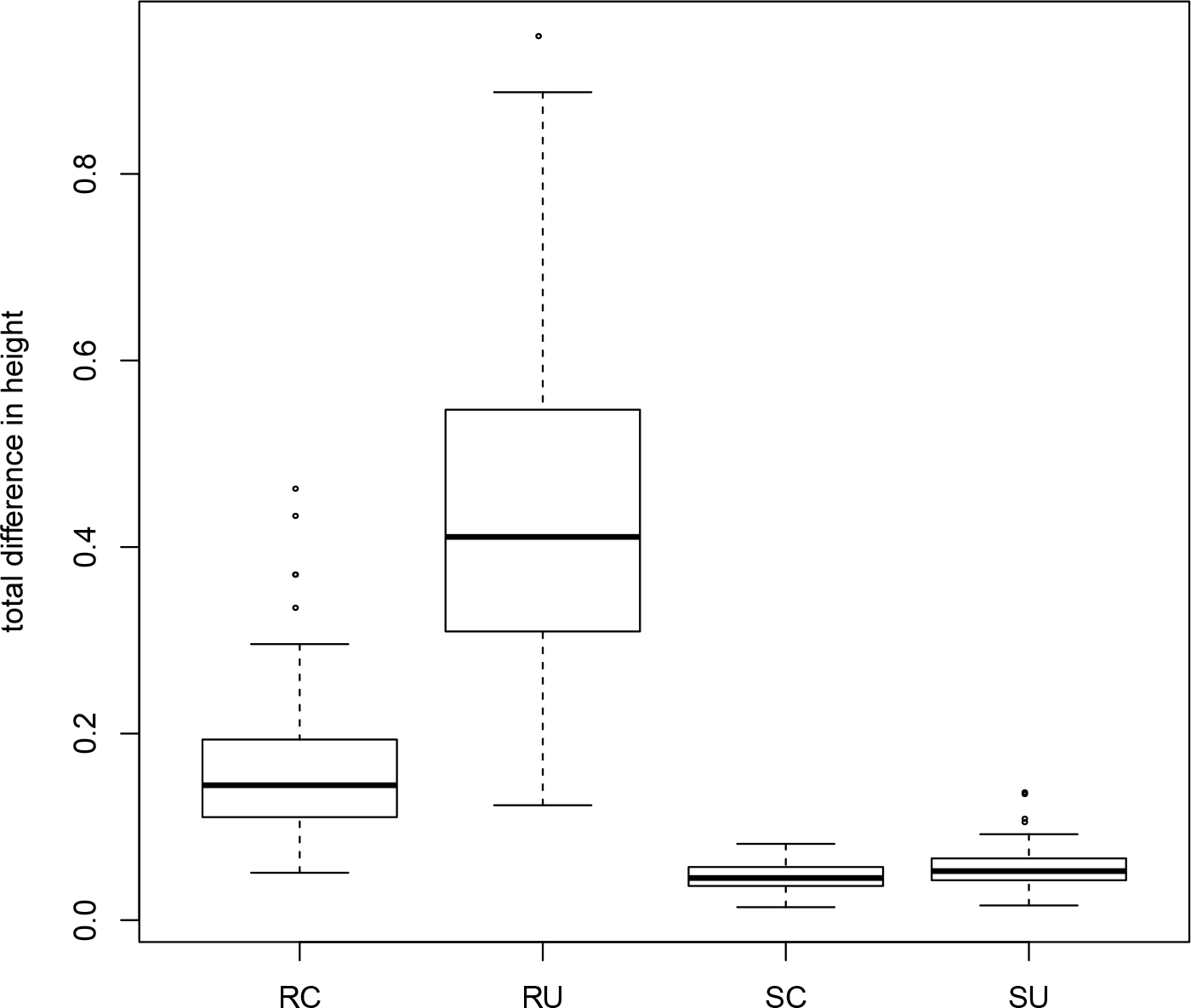
Results from a simulation comparing total difference between true and estimated ages for three random genes under a clock (RC) and UCLN (RU) and three genes chosen with SortaDate under a clock (SC) and UCLN (SU). Results are from 100 simulations of distinct parameter values. See details in the text.

### Reconsidering standard clock tests

Several gene trees from the examples discussed fail a standard strict clock test but have low root-to-tip variance. To briefly explore whether this was the result of the stringency of the clock test, we simulated strict clock amino acid and nucleotide data on orthologs from each empirical dataset and examined the frequency of incorrectly rejecting a strict clock. The false rejection rate for clock tests using amino acid data and a strict clock were between 5% and 8%. For the two nucleotide datasets, the rejection rate was much higher at 23% and 46%. This suggests that for amino acid data, the false rejection rate was near the nominal value, while for the nucleotide datasets the false rejection rate was unreliable. Both nucleotide datasets (BIR and CARY) also had the largest number of species and so the rejection rate may be a function of the number of taxa (i.e., with a greater number of sampled lineages, cumulative stochastic variation for even clock-like data can lead to the rejection of a strict clock). Sensitivity of the clock-test to nucleotide data is not the focus of this study, but should be examined in more detail. Furthermore, it would be more informative to examine the deviation from the clock instead of a boolean test of significant fit. In regard to divergence-time estimation, if a strict or stricter clock can be used, molecular phylogenies may be dated with significantly lower error. As an added benefit, fewer fossils would be necessary to calibrate nodes (and indirectly, rates). We suggest that the community explore model fit to relaxed clock models as well as potential alternatives to the prevailing strict clock test that may be more beneficial for divergence-time estimates and more informative in regard to rate heterogeneity in phylogenomic datasets.

## Acknowledgements

We would like to thank Oscar Vargas, Greg Stull, Jianjun Jin, Drew Larson, Ning Wang, Caroline Parins-Fukuchi, Edwige Moyroud, Lijun Zhao, and Nat Walker-Hale for discussion of the manuscript and method. Greg Stull assisted with testing the SortaDate code. JFW and SAS were supported by NSF 1354048. JWB and SAS were supported by NSF 1207915.

## Data availability

While the data used here was all public, we conducted some orthology analyses, the results of which are available on GitHub at. Associated scripts related to the method are available on GitHub at https://github.com/FePhyFoFum/sortadate.

## Author contributions

SAS, JWB, and JFW conceived of the project. SAS, JFW, and JWB analyzed the data. SAS, JWB, and JFW wrote the manuscript.

